# LSD Restores Synaptic Plasticity in VTA of Morphine-Treated Mice and Disrupts Morphine-Conditioned Place Preference

**DOI:** 10.1101/2025.06.12.658958

**Authors:** Michael Von Gunten, Tenna Russell, Isaac Stirland, Seth Parks, Timothy Jenkins, Jeffrey G. Edwards

## Abstract

Psychedelics are emerging as a promising treatment option for a range of neuropsychiatric disorders, including substance use disorders. One potential mechanism underlying their therapeutic benefits may involve a reversal of maladaptive plasticity induced by drug exposure. Here, we identify physiological, behavioral, and epigenetic impacts of lysergic acid diethylamide (LSD) on morphine-treated male and female mice. Morphine was selected due to the high leverage capacity to address the opioid epidemic. A single treatment of LSD, or 4 microdoses of LSD, cause accelerated extinction of morphine-induced conditioned place preference. Whole-cell electrophysiology revealed that excitatory synaptic plasticity, which was eliminated in VTA GABA neurons following morphine exposure, was restored 24 hours after a single high dose of LSD. To explore the impact of LSD treatment on potential epigenetic changes, whole-brain DNA methylation analysis in morphine-treated mice that received either saline or LSD post-morphine treatment revealed significant differences in methylation profiles associated with LSD treatment. Collectively, these findings suggest that LSD may reverse or prevent morphine-induced changes in reward circuit plasticity and attenuate measures of morphine-preference.

## Introduction

In 2022 there were over 70,000 opioid overdose deaths in the united states [1]. While opioids are commonly prescribed as analgesics, their high addictive potential poses a severe public health crisis known colloquially as the opioid epidemic. These drugs modulate the mesolimbic system by acting on mu opioid receptors (MORs) [2], which suppress presynaptic GABA release onto the DA neurons. This disinhibition increased dopamine (DA) release from the ventral tegmental area (VTA), thereby mediating reward signaling [2, 3]. Although opioids acutely induce reward, long-term opioid exposure alters synaptic plasticity in both VTA GABA and DA neurons [4, 5], contributing to drug craving and withdrawal symptoms [6, 7]. Despite extensive research examining treatment options for opioid use disorder (OUD), no existing treatment directly targets this maladaptive plasticity in the brain’s reward system.

Current treatments for OUD include inpatient and outpatient therapy, or administration of alternative opiates that target MORs. Unfortunately, clinicians report extremely high relapse rates with these methods [8, 9], prompting researchers to explore alternative therapies, including the use of classic psychedelics such as lysergic acid diethylamide (LSD) and psilocybin to treat OUD [10] and other substance use disorders (SUDs) [11–13]. Psychedelic-based treatments come in two main forms: short-term (1-3 treatments) high dose administration which is inherently hallucinogenic [14, 15], or a microdosing approach where the acute sensory altering aspects of the drug are largely imperceptible [16–18]. Both treatments show promise in treating various mental disorders including SUDs [12, 18–21], although the mechanisms underlying their efficacy remain a topic of debate [22, 23].

One proposed mechanism suggests that psychedelics promote structural and functional plasticity [24–27] via their actions on serotonin-2A (5-HT2A) or tropomyosin receptor kinase B (TrkB)-receptors [27, 28]. For instance, LSD treatment leads to increased dendritic spine density and arborization in rodent cortical neurons [27], and enhances α-amino-3-hydroxy-5-methyl-4- isoxazole propionate (AMPA) receptor-mediated currents in the PFC [29]. Notably, 5-HT2A receptors are expressed by the DA and GABA neurons in the VTA [30, 31], and are also highly expressed in the PFC [32] which sends glutamatergic innervations to the VTA. While plasticity may play a key role in psychedelic treatment of mental disorders, other researchers propose that alterations to DNA methylation mediate their therapeutic effects [33]. To further elucidate the biological mechanisms underlying the use of psychedelics in treating substance use disorders (SUDs), our study investigates the behavioral, physiological, and epigenetic effects of either a single high dose or multiple low doses of LSD on morphine-treated mice.

## Methods

All experiments were performed in accordance with Institutional Animal Care and Use Committee protocols and followed National Institute of Health Guide for the care and use of laboratory animals. Institutional Animal Care and Use Committee protocols for all experiments were approved by the Brigham Young University Institutional Animal Care and Use Committee, Animal Welfare Assurance Number A3783-01.

### Animals

Male and female CD1 GAD67-GFP knock-in mice (heterozygous; P21–P65) produced by the Tamamaki laboratory were used to fluorescently identify VTA GABA cells [34]. In a prior report, we confirmed via PCR that extracted GFP-positive cells, but not DA cells, express GAD67 and performed immunohistochemistry experiments that confirm TH and GFP-positive cells are separate populations in the VTA [35]. GFP-positive mice were identified initially in screening using UV goggles.

### Drug treatments

Mice were assigned to one of three treatment groups and received intraperitoneal (IP) injections according to the designated schedule. One group received morphine (10 mg/kg) every other day on days 0, 2, 4, and 6, with saline injections on alternating days (days 1, 3, 5, and 7). A second group received the same morphine treatment schedule but was co-administered a low dose of LSD (0.03 mg/kg) alongside morphine on days 0, 2, 4, and 6, while receiving saline on the alternating days. The third group also received morphine on days 0, 2, 4, and 6, with saline on days 1, 3, and 5; however, instead of saline on day 7, these mice received a high dose of LSD (0.3 mg/kg). For the conditioned place preference experiments, the single high dose of LSD (0.3 mg/kg) was administered directly after conditioning. The high dose of LSD (0.3 mg/kg) was chosen based on behavioral effects seen by Winter et al [36] and Rodriguiz et al [37]. For the low dose of LSD, we based our dose off of work from De Gregorio et al [29] who noted that behavioral effects were not seen after a single injection of 0.03 mg/kg LSD but were seen after repeated treatments, suggesting it as an approximation of sub-hallucinogenic dose in humans. The rationale for these two concentrations is to correlate our work in rodents with studies in humans that often employ high or low doses reflecting either hallucinogenic or non-hallucinogenic concentrations, respectively [39–41]. Whole-cell experiments were performed 24 hours after the last injection.

### Slice preparation

Mice were anesthetized with isoflurane and decapitated with a rodent guillotine. Directly prior to brain extraction, cardiac perfusions were performed using 10 mL ice cold, oxygenated NMDG cutting solution that was injected into the left ventricle of the mouse’s heart. After the cardiac perfusion brains were rapidly removed and sectioned horizontally on a vibratome at 250 μm using an ice-cold N-methyl-D-glucamine (NMDG)-based cutting solution composed of 92 mM NMDG, 2.5 mM KCl, 1.25 mM NaH_2_PO_4_, 30 mM NaHCO_3_, 20 mM HEPES, 25 mM glucose, 2 mM thiourea, 5 mM Na-ascorbate, 3 mM Na-pyruvate, 0.5 mM CaCl_2_·2H_2_O, and 10 mM MgSO_4_·7H_2_O, pH balanced to 7.5 pH with 12 M HCL. After sectioning, adult slices were kept in a warm (34◦C) bath of the cutting solution and were spiked with increasing amounts of NaCl over a 10 min period as described by Ting et al [42] before being placed in an oxygenated HEPES holding solution composed of 92 mM NaCl, 2.5 mM KCl, 1.25 mMNaH_2_PO_4_, 30 mM NaHCO_3_, 20 mM HEPES, 25 mM glucose, 2 mM thiourea, 5 mM Na-ascorbate, 3 mM Na-pyruvate, 2 mM CaCl_2_·2H_2_O, and 2 mM MgSO_4_·7H_2_O pH balanced to 7.5 with 10 M NaOH.

### Recording protocol experimental design

Recordings began at least 1 h after brain extraction and slicing. Slices were placed in the recording chamber and bathed with oxygenated (95% O_2_, and 5% CO_2_) high divalent ACSF consisting of 119 mM NaCl, 26 mM NaHCO_3_, 2.5mM KCl, 1 mM NaH_2_PO_4_, 4 mm CaCl_2_, 4 mm MgCl_2_ and 11mM glucose, with 100 μm picrotoxin to block GABA_A_ currents. The VTA was visualized using an Olympus BX51W1 microscope with a 40× water-immersion objective. Cells were selected at approximately the following coordinates from adult mouse bregma; anteroposterior −3.2 to 3.3, mediolateral ±0.6 to 0.7, dorsoventral −4.5 to 4.3 and identified via GFP fluorescence. Patch pipette resistance was 2.5–5.5 mΩ. GABA cells were held at −65 mV and patched with a glass pipette filled with internal solution composed of 117 mm cesium gluconate, 2.8 mm NaCl, 20 mm HEPES, 5 mm MgCl_2_, 2.17 mM ATP-Na, 0.32 mM GTP-Na and 1 mm QX-314 at pH 7.28 (275–285 mOsm). Current traces were recorded using Multiclamp 700B amplifier (Molecular Devices). Signals were filtered at 4 kHz and digitized with an Axon 1550A digitizer (Molecular Devices) connected to a Dell personal computer with pClamp 10.5 Clampex software (Molecular Devices). The peak amplitude of the induced EPSC was calculated using Clampfit 10.7 software (Molecular Devices) and graphed using Origin 8. Within individual experiments, current amplitudes were averaged by minute (6 sweeps/min). Plasticity was induced using two stimulations at 100 Hz for 1 s each, separated by a 20 second interval (high frequency stimulation; HFS).

### Statistical analysis

To determine if significant LTD had occurred within the same experiment (each individual cell) a one-way ANOVA was used within each treatment group comparing the 0-5 min before conditioning to 10-15- and 15-20-min post-conditioning. To compare post-HFS EPSCs across treatment groups, a one-way ANOVA with a Tukey post-hoc analysis was used. N values signify the number of cells used in each experiment, where 1–2 cells were acquired per animal (minimum of 4 biological replicates per treatment group). To assess statical significance the alpha level was set to 0.05.

### Conditioned place preference

Our conditioned place preference protocol was adapted from Cunningham, et. al., 2006 [43]. The conditioned place preference (CPP) apparatus was composed of 3 compartments, a neutral middle chamber, a chamber with visual cues on the walls and a dimpled texture floor, and a chamber with no visual cues on the walls and a line-textured floor.

The three-compartment apparatus was selected to remove the potential bias caused by a two-compartment apparatus [44]. The dimensions of the drug- or vehicle-paired chambers were 20 cm x 20 cm x 18 cm, and the dimensions of the middle chamber were 20 cm x 6.5 cm x 18 cm. First, mice were allowed to habituate to the CPP chambers for two days, 30 minutes each day. This was followed by a pretest to assess initial preference to either chamber (bias design).

Similar to other studies, we found that mice reliably develop morphine-induced CPP using a biased design [45], therefore we employed this model. After habituation and initial preference testing, mice were subjected to 4 drug (morphine (10mg/kg) or morphine (10mg/kg) combined with a low dose of LSD (0.03mg/kg)) conditioning sessions in the less preferred chamber on days 0, 2, 4, and 6. On days 1, 3, 5 and 7, saline (vehicle control) conditioning sessions were conducted in the more preferred chamber (see Figure 1). Each conditioning session lasted for 50 minutes. Preference testing was performed for the subsequent 3 days after the last conditioning session. Testing sessions lasted for 30 minutes. Conditioning and testing sessions were performed between 3-7 pm. Behavior was videoed with a GoPro camera and Anymaze data analysis software was used to measure the amount of time spent in each chamber and distance traveled during testing. CPP scores were calculated with the following formula, (percent time CS^+^_post_ - percent time in CS^+^_pre_) / percent time CS^+^_pre_) *100 [46]. The researchers conducting the experiments were blinded to the treatment being administered.

**Figure 1.**
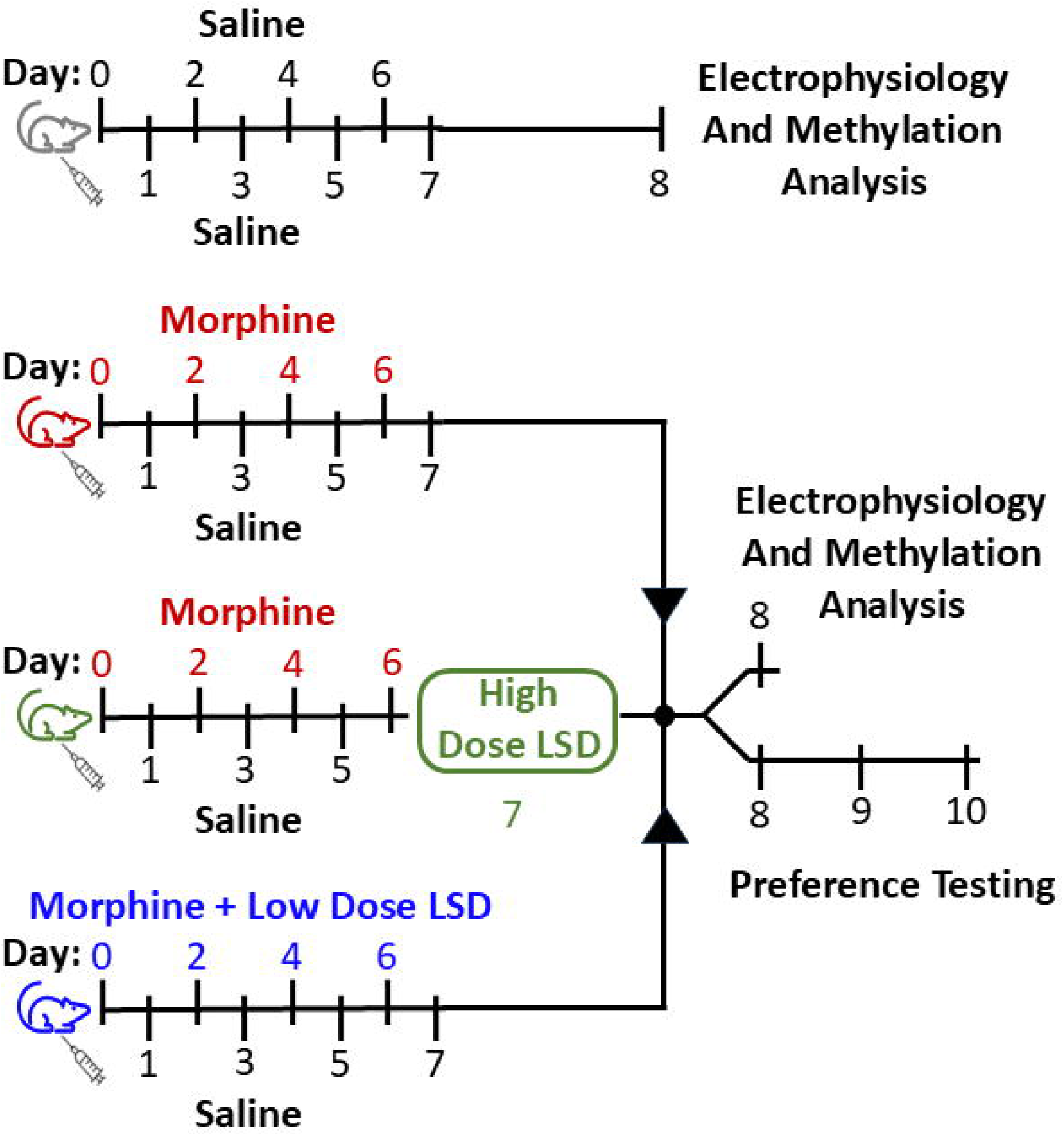
Experimental design. Timeline depicting the treatment schedule for the four experimental cohorts.

### DNA Methylation

Whole brain tissue was mechanically homogenized prior to DNA isolation using the DNeasy Blood and Tissue Kit (Qiagen, Germantown, Maryland). DNA was then bisulfite converted using the EZ DNA Methylation Kit (Zymo Research, Irvine, California). Next, methylation analysis was performed using the Infinium Mouse Methylation BeadChip array and was processed at the University of Utah Genomics Core Facility. All analyses of array data were conducted using R Statistical Software (v4.4.0; GUI 2023). Using the minfi package, array data in the form of idat files were converted into fraction methylation (beta) values, annotated with the mm10 genome, and normalized (SWAN). A sliding window approach using the USEQ software [47] was implemented to identify differentially methylated regions (DMRs). Only regions with a Phred-Scaled FDR ≥ 13 were considered significant. All subsequent analyses were conducted using base R commands and the car package. The Stanford GREAT online tool and the UCSC Genome Browser were implemented for ontological assessment of identified DMRs [48].

## Results

### Morphine treatment induces conditioned place preference

The methodological approach timeline is outlined in Figure 1. Initially we established that mice show morphine preference in our conditioned place preference (CPP) assay. Mice were treated with morphine (10 mg/kg) and preference was examined using a biased conditioned place preference paradigm. Morphine-treated mice manifested preference for the morphine-conditioned chamber that did not diminish over a three-day period post-conditioning (Figure 2A).

**Figure 2:**
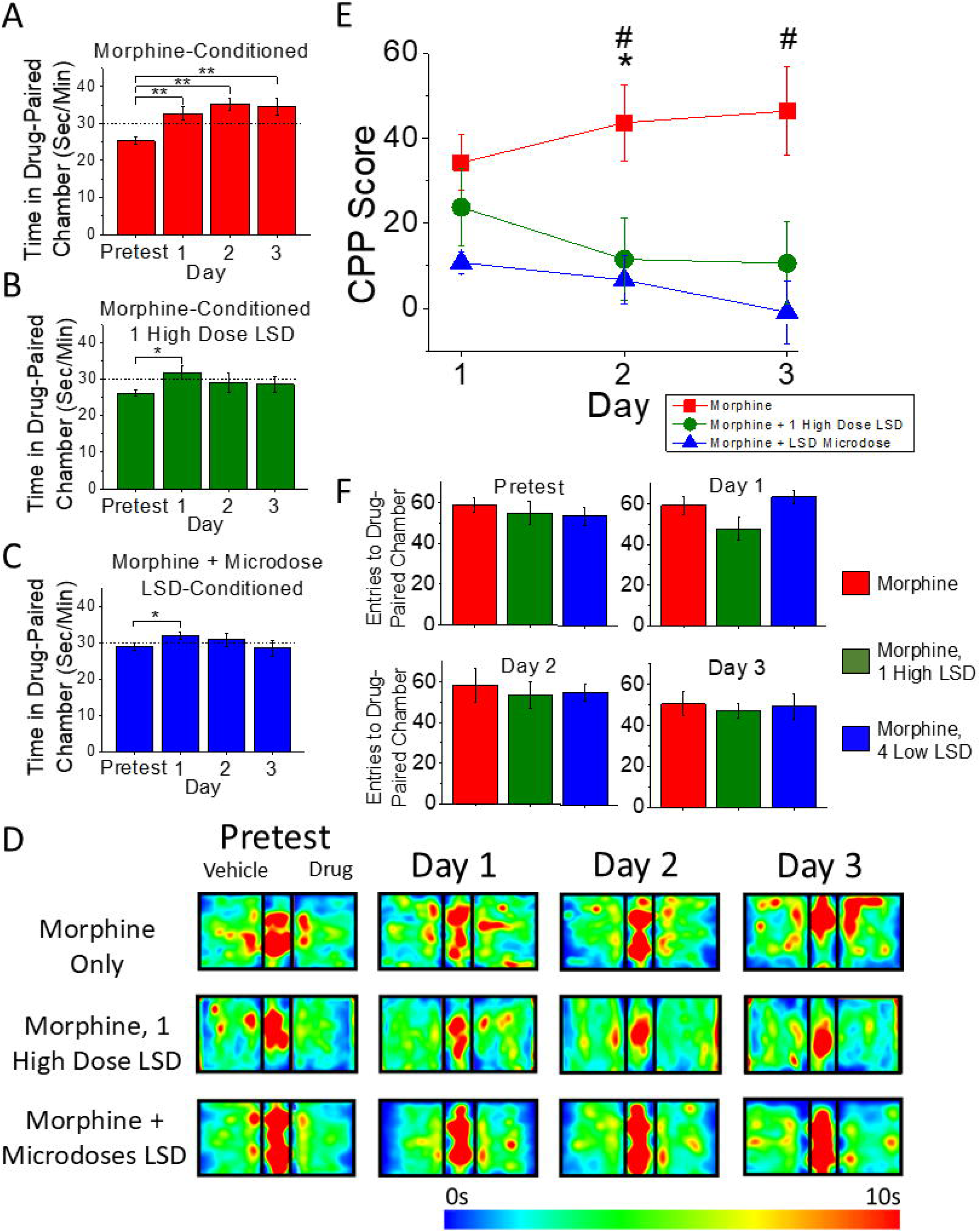
A high dose of LSD post morphine-conditioning, or 4 low doses of LSD during morphine-conditioning, both lead to accelerated extinction of morphine preference. **A)** Morphine-conditioned mice spent significantly more time in the morphine-paired chamber compared to the pretest for 3 days post-conditioning (n = 10, p < 0.01 for all 3 days). **B)** Morphine-conditioned mice who received a single high dose of LSD after the morphine-conditioning regimen spend significantly more time in the morphine-paired chamber on the first day of preference testing (n = 9, p < 0.05) but did not spend significantly more time on days 2 and 3 (p > 0.05). **C)** Morphine and low dose LSD-conditioned mice spend significantly more time in the morphine + LSD-paired chamber on the first day of testing (n = 9, p < 0.05) but did not spend significantly more time on days 2 and 3 of preference testing (p > 0.05). **D)** Comparison of CPP scores between each of the treatment groups shows no significant difference in CPP scores on the first day of preference testing (p > 0.05), and significantly decreased CPP scores on days 2 and 3 of preference testing for both the single high dose of LSD, and the 4 low doses of LSD groups compared to the morphine-treated mice who did not receive any LSD treatment. (p < 0.05). * indicates significant difference between morphine-conditioned mice and morphine-conditioned mice that received a high dose of LSD (0.3 ug/kg). # indicates significant difference between morphine-conditioned mice and morphine + micro-LSD-conditioned mice. **E)** The number of entries into the drug paired compartment was not different across any of the groups for any of the days (p > 0.05). **F)** Average heat plots for the pretest day, and each of the preference testing days for each of the treatment groups. *Error bars represent standard error of the mean*.

### LSD treatment accelerates extinction of morphine-preference

To assess the effect of LSD-treatment on morphine preference we examined two different concentrations of LSD. To test the effects of a high dose of LSD on morphine-treatment, morphine-conditioned mice received a single high dose of LSD (0.3 mg/kg) or saline as a vehicle control. The morphine-conditioned mice who received a saline treatment manifested preference for the morphine-pared chamber for 3 days post-conditioning (Figure 2A). Notably, the morphine-conditioned mice who were treated with a high dose of LSD showed preference for the morphine conditioned chamber during the first day of preference testing, but no preference was observed during the second and third days of testing (Figure 2B). Because LSD microdosing may also yield therapeutic benefits, we conditioned a cohort of mice to a combination of morphine (10mg/kg) and a 10-fold lower dose of LSD (0.03mg/kg; see Figure 1 injection protocol). These mice exhibited a pattern similar to mice treated with a single high dose of LSD, showing preference during the first day of treatment, but showing no significant preference during the subsequent two days (Figure 2C, see also Figure 2D). Further analysis assessing CPP scores across the treatment groups shows significantly decreased CPP scores for mice treated with a single high dose of LSD and the mice treated with microdoses of LSD compared to the mice treated with only morphine (Figure 2E). No significant differences in CPP score were noted between the mice treated with 1 high dose of LSD after morphine conditioning and the mice treated with low doses of LSD during morphine conditioning (Figure 2E). Number of entries into the drug paired chamber was not significantly different between groups (Figure 2F).

### Morphine treatment eliminates LTD

Drug-induced alteration of excitatory plasticity in the VTA is a well-established cellular hallmark of drug dependence. Therefore, we next examined the impact of LSD on morphine-induced plasticity changes. To measure these plasticity alterations, we examined excitatory LTD of VTA GABA cells using whole cell electrophysiology. Mirroring the behavioral experiments, mice were treated with four doses of morphine (10 mg/kg), each administered 48 hours apart. Littermates of these mice were treated with saline (vehicle control). After the treatment regimen, whole cell electrophysiology was performed on VTA GABA cells and EPSC amplitude was recorded. VTA GABA cells of saline-treated mice manifested typical LTD (Figure 3A) as we have noted previously [49, 50], however LTD was abolished in the morphine-treated mice (Figure 3B).

**Figure 3:**
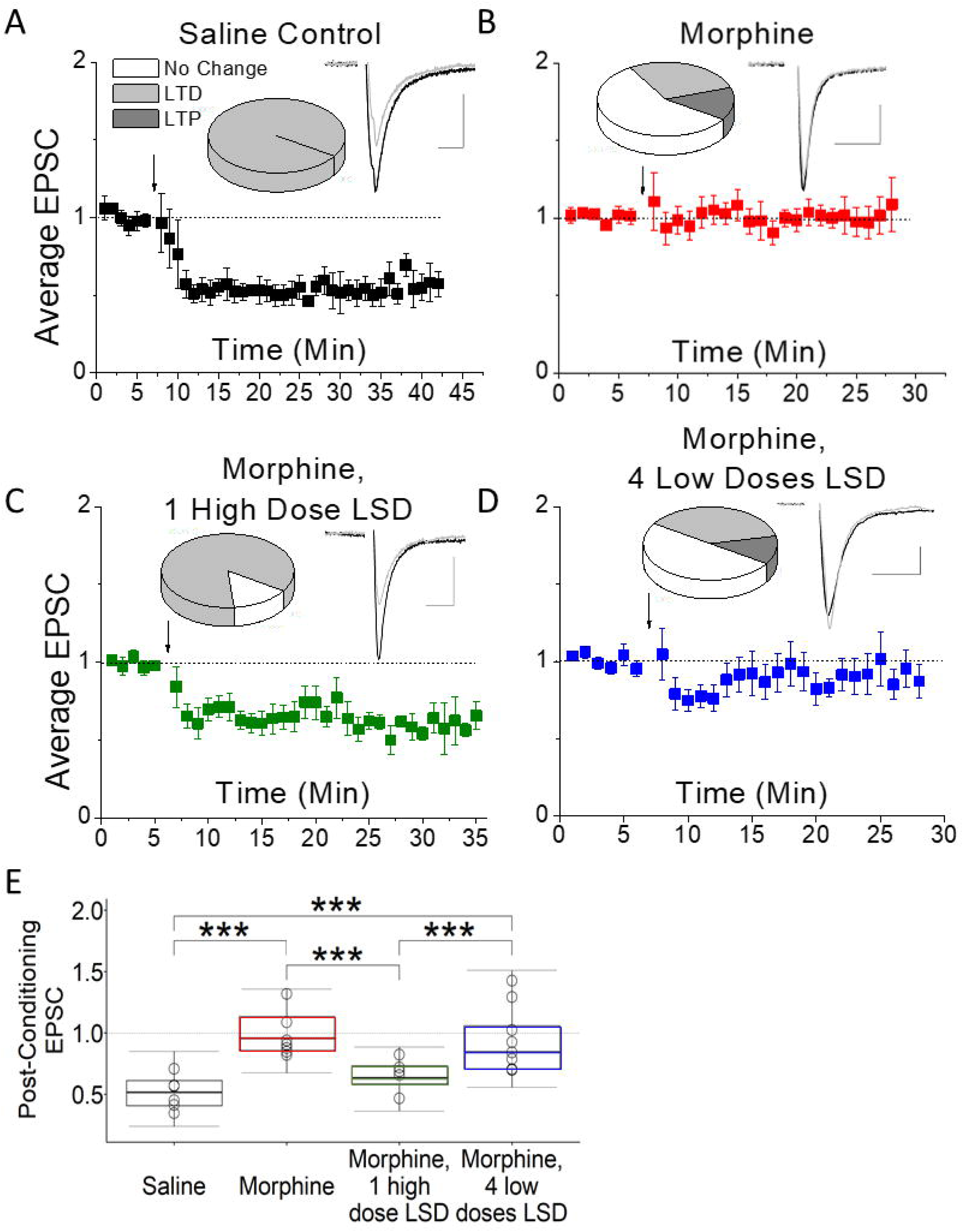
A high dose of LSD restores excitatory LTD in VTA GABA cells of morphine-treated mice. **A)** High frequency stimulation (HFS; arrow) induces long-term depression (LTD) in mice treated with saline (vehicle) injections 1/day for 8 days (n = 6 cells from n = 4 mice; p < 0.001. at both 10-15 and 15-20 min post HFS). **B)** Four morphine injections administered every 48 hours, with four saline injections given on the days between morphine treatments, eliminates LTD (n = 9 cells from n = 5 mice; p = 0.82 10-15 min post HFS and p = 0.95 16-20 min post HFS). **C)** LTD is restored in morphine-treated mice 24 hours after 1 injection with LSD (0.3mg/kg; n = 7 cells from n = 4 mice; p < 0.001 at both 10-15 and 15-20 min post HFS). **D)** Four treatments of combined morphine (10mg/kg) and LSD (0.03mg/kg;) with 4 saline injections on alternating days eliminates LTD (n = 8 cells from n = 5 mice; p > 0.05 at 10-15 min post HFS and p > 0.05 at 15-20 min post HFS. **E)** Further analysis of post-HFS EPSC amplitude was done using an ANOVA with a Tukey post-hoc test to compare between treatment groups. This analysis revealed that the post-HFS EPSC amplitude for morphine-treated mice at 10-15 min post-HFS was significantly different from the saline-treated mice (p < 0.001), and from the morphine treated mice that received 1 high dose of LSD (p < 0.001) but was not different from the mice who received the combined morphine and low dose of LSD treatment (p > 0.5). This analysis also showed no significant difference between the saline-treated mice and the morphine-treated mice who received 1 high dose of LSD (p > 0.05). *Glutamate currents were recorded from GAD67-GFP-positive VTA GABA cells. All plots are normalized EPSC amplitude means with standard error of the mean. Arrow indicates 100 Hz HFS conditioning induction. Insets, here and throughout, are representative averaged EPSC traces (10–12 traces) taken from before (black) and 10–15 min after conditioning (gray). Calibration: Scale bars are 50 pA, 5 ms*.

### A single high dose of LSD restores morphine-eliminated LTD

To assess the impact of LSD treatment on morphine-induced elimination of LTD in VTA GABA cells, morphine-treated mice received one high dose of LSD (0.3mg/kg) twenty-four hours after their final morphine treatment. Twenty-four hours following LSD administration we performed whole-cell electrophysiology. Interestingly, excitatory LTD was restored in the VTA GABA cells of mice treated with a high dose of LSD (Figure 3C). In contrast, co-administration of a low dose of LSD (0.03mg/kg) with morphine (10mg/kg), failed to prevent elimination of LTD (Figure 3D). Further analysis across treatment groups revealed that the magnitude of LTD recorded in the high dose LSD-treated mice was not significantly different from the LTD recorded in saline-treated mice (Figure 3E).

### LSD treatment causes differential DNA methylation compared to control and morphine groups

Due to the interesting electrophysiological results observed in the mice treated with morphine and a high dose of LSD (Figure 3C), we elected to analyze brain methylation patterns in mice treated with a saline control, morphine, or morphine and a high dose of LSD. Three sliding window analyses were implemented to iteratively compare control, morphine-treated, and morphine-treated and a high dose of LSD groups to one another. These analyses identified two regions of differential methylation, which mapped to the same genomic locus (chr7:13261083-13261446). This region was identified as significantly differentially methylated (Phred-Scaled FDR >13) between the control group and the morphine-treated group that received the high dose of LSD, as well as between the morphine-treated group and the morphine-treated group who received a high dose of LSD (Figure 4A). This suggests that methylation profiles between LSD treatment is associated with a significant change to methylation respective to the control and morphine-treated groups. The identified region of differential methylation (chr7:13261083-13261446) was input to the UCSC Genome Browser, which revealed its location within the coding region for Zswim9.

**Figure 4.**
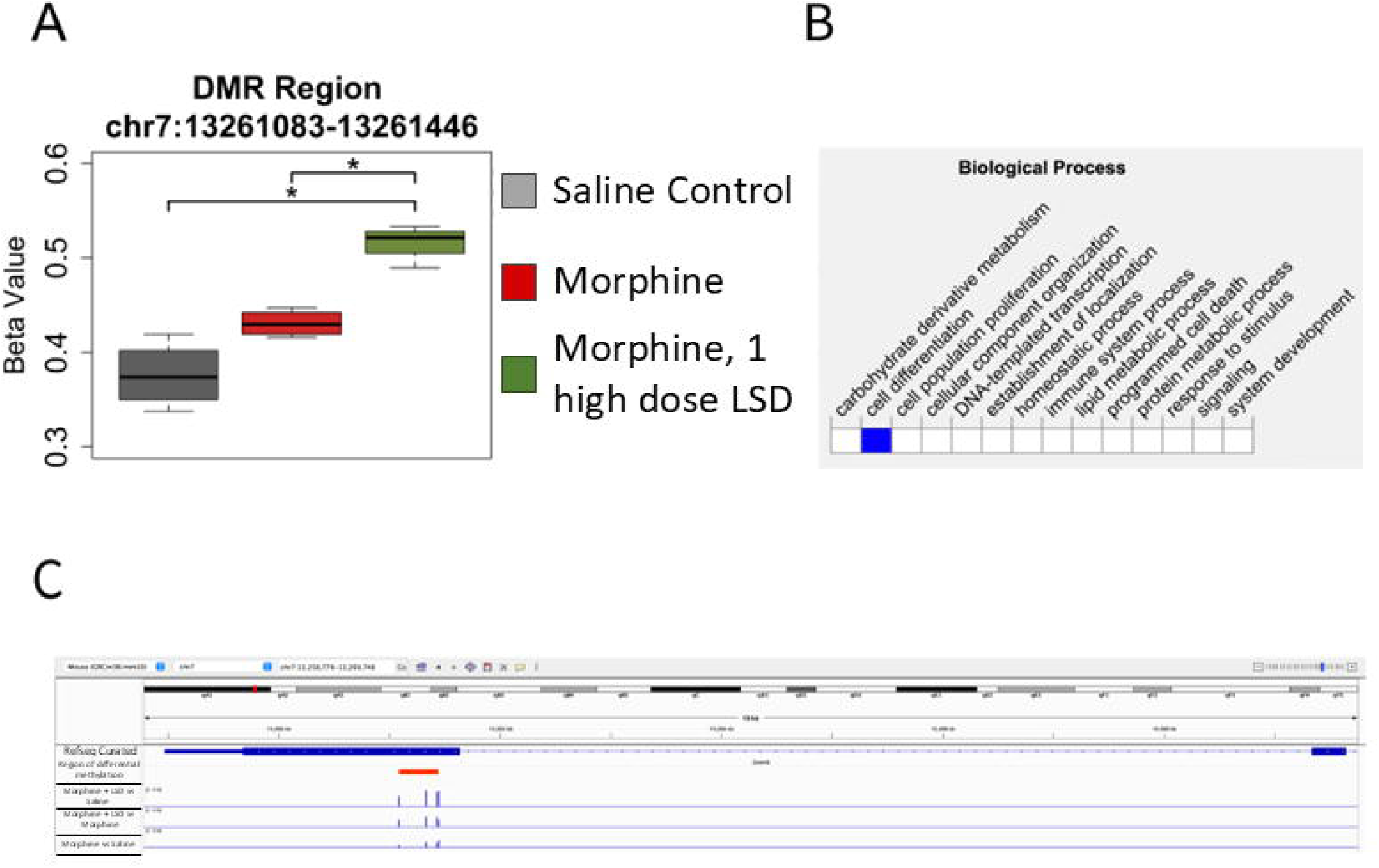
A high dose of LSD alters DNA methylation in morphine-treated mice. **A)** Differences in DNA methylation levels at a specific region (p < 0.05, Phred-Scaled FDR > 14) identified by the USEQ sliding window analysis between Saline Control, Morphine, and Morphine, 1 High Dose LSD sample groups. **B)** Screenshot from Mouse Genome Informatics (MGI) of the oncologic analysis for the genomic region identified as differentially methylated amongst groups. **C)** Integrated Genomics Viewer (IGV) screenshot of the differentially methylated region located within an exon of the Zswim9 gene. The first (top) track is the RefSeq genome. The second highlights the region of differential methylation (363bp). The third, fourth and fifth tracks show the magnitude of differential methylation, at each CpG within the differentially methylated region, between the multiple sample groups and are scaled to the same height. Track three displays the difference in methylation between the morphine + LSD group and the saline control group. Track four displays the difference in methylation between the morphine + LSD group and the morphine group. Track five displays the difference in methylation between the morphine and control groups.

## Discussion

The demonstration that LSD reverses morphine-induced changes to synaptic plasticity, attenuates morphine-induced place-preference, and alters DNA methylation patterns in morphine-treated mice has potential clinical applications. The is extremely provocative data as it relates to potential cellular correlations of the capacity for LSD to treat opioid related disorders.

### LSD impact on morphine-preference

We first demonstrated the reliability of our model as mice treated with morphine prefer the morphine paired chamber for three days post-conditioning, consistent with previous reports [51–53]. Notably, mice treated with a single high dose of LSD (0.3mg/kg) 24-hours before preference testing only show morphine preference for 1-day post-conditioning and manifest significantly lower CPP scores on days two and three post conditioning. Psychedelics impact on behavior is likely through 5-HT2a receptor activation [54], which attenuates aversive responses to drug withdrawal [55, 56]. Furthermore, similar experiments assessing morphine CPP in mice were performed after a high dose of ketamine and showed decreased morphine preference as soon as 30 minutes after ketamine treatment [45].

In addition, our mice conditioned to morphine combined with a low dose of LSD (0.03mg/kg) also showed morphine preference during the first day of preference testing but did not show any morphine preference on days two or three. In this case, it should be considered that extinction of drug preference could be due to the acute effects of the low dose of LSD during the morphine conditioning trial, in addition to LSD preventing some of the long-term neurological effects of morphine. Future experiments should be done in which the low dose of LSD is administered after each morphine-conditioning trial, as this would parse out any acute effects of the low dose of LSD during the morphine conditioning trials. Collectively, these findings illustrate that psychedelic treatment can modulate opioid-induced behavioral alterations in mice, supporting the growing body of evidence that LSD may hold promise as a treatment option for OUD.

### LSD restores excitatory synaptic plasticity impaired by morphine

Glutamatergic plasticity in the VTA is necessary for rodents to manifest morphine-induced conditioned place preference [57], and addictive psychoactive substances altering VTA glutamate plasticity is a hallmark of drug dependence creation [6]. Therefore, we sought to understand how morphine-induced alterations to VTA glutamatergic plasticity correlate with our findings from the conditioned place preference experiments, and whether psychedelic treatment can reverse morphine-induced plasticity changes. LTD of VTA GABA cells was an excellent assay system as we determined morphine exposure abolishes this LTD. Thus, preventing or reversing mesocorticolimbic opiate-induced synaptic changes may be central for successful pharmacological treatment of OUD.

Although plasticity alterations play a key role in OUD, the potential for psychedelic treatment to reverse these changes in the VTA remains largely unexplored, underscoring the relevance of the present study. Interestingly, we note that a single high dose of LSD restores LTD in morphine-treated mice. This restoration of plasticity in the VTA is key to understanding the mechanism through which psychedelics may be working to treat drug dependence and other neurological disorders. This is the first report, to our knowledge, providing electrophysiological evidence that LSD can reverse maladaptive plasticity in the mesolimbic system, an essential brain pathway involved in the development of drug dependence. Although studies are limited, a few have examined psychedelic impact on plasticity in other brain regions, with one demonstrating enhanced long-term potentiation (LTP) in the prefrontal cortex [58]. These findings support out current study in the reward circuit by providing precedent for psychedelic-induced plasticity changes within an area that sends excitatory inputs to VTA GABA cells [59].

There is mounting evidence demonstrating that psychedelics such as LSD induce robust structural and functional plasticity [25, 27, 60], through targeting 5HT2A and TrkB receptors [28, 61, 62]. It is postulated that this plasticity is a main mechanism through which psychedelics work to treat neurological disorders [25, 60]. In fact, maladaptive plasticity is one of the few commonalities between the diverse array of neurological disorders that psychedelics are used to treat, ranging from PTSD [63–65] and SUDs [66], to seemingly unrelated disorders such as cluster headache [67, 68]. Researchers have proposed that psychedelics work to reset maladaptive brain circuitry [69], or to reopened critical periods [70], and a restoration of plasticity could underlie either of these theories.

We propose that the restoration of LTD in VTA GABA neurons following LSD administration may be mediated by the formation of new synapses or by functional recovery of previously impaired synapses. Indeed, psychedelics are reported to induce structural plasticity and synaptic outgrowth that are maintained for up to one month even after a single exposure [27, 71], and it is plausible that newly formed synapses, naïve to morphine’s effects, may underlie this re-emergence of LTD. However, it is also possible that LSD acts on pre-existing synapses, restoring their ability to undergo plasticity after being impaired by prior morphine exposure.

Future studies should examine these potential causes, along with plasticity restoration in other brains regions, or induced by other events such as chronic stress [72].

Although mice treated with low-dose LSD during morphine conditioning exhibited reduced morphine preference behaviorally, LTD was not recovered. A possible explanation for this is that while measuring excitatory LTD to VTA GABA cells is one form of synaptic plasticity characterization, other circuits could also be impacted and further examined (i.e. VTA DA cell plasticity) that mediate low-dose LSD behavioral changes. Nevertheless, the behavioral data strongly suggest that low-dose LSD mitigates morphine-induced preference in mice, even without fully defining the underlying circuit mechanisms. This aligns with broader findings that psychedelic microdosing can reduce behavioral anxiety [73, 74], supporting its potential as a therapeutic strategy for preventing opioid dependence.

### Morphine and LSD impact on whole-brain DNA Methylation

Drugs of abuse are thought to induce dependence through a combination of factors including modifications to cellular activity, synaptic plasticity, and gene transcription via epigenetic mechanisms such as DNA methylation and acetylation [75, 76]. Importantly, therapeutic effects of psychedelics likely involve DNA methylation [33]. Indeed, a single LSD dose (0.2 mg/kg) altered gene expression patterns in rats [77]. Therefore, we sought to investigate whether LSD can modify epigenetic signatures in the brain after morphine treatment by examining DNA methylation patterns.

Among the loci analyzed in our study, Zswim9 emerged as a region with significant differential methylation in the morphine-treated mice that received a high dose of LSD. Members of the Zswim family (Zswim1-9) are zinc finger proteins expressed in many tissues and specifically are highly expressed in the brain during development [78–80]. Zinc finger genes are DNA recognition proteins that are associated with psychosis-related disorders, particularly within dopaminergic pathways [81, 82]. Additionally, zinc plays a role in glutamatergic neurotransmission via its association with glutamatergic synaptic vesicles, linking zinc-related proteins with other forms of neuropathology [83]. Zswim9 is therefore well situated to be involved in potential epigenetic changes that are relevant to reward signaling. However, while patterns of DNA methylation often associate with differential gene expression, establishing a causal relationship to phenotypic outcomes remains complex, though numerous reports highlight weak relationship between differential methylation and phenotype [84–86].

While the functional consequences of differential methylation remain to be fully elucidated, the identification of a single locus at the Zswim9 gene that was differentially methylated in morphine-exposed mice after a high dose of LSD suggests a specific epigenetic signature associated with LSD treatment. When considered alongside our electrophysiological and behavioral findings, these data support the concept that LSD alterations to DNA methylation may be a potential functional target of LSD. Further studies, including RNA-sequencing and gene expression profiling, are warranted to determine the downstream functional relevance of this epigenetic modification.

## Conclusion

Our data reveals LSD reverses plasticity and behavior impact of morphine in mice, while also inducing epigenetic modifications that together have the potential for understanding the broader impact of psychedelic treatment. Collectively, these findings provide preclinical evidence highlighting the potential for LSD as a possible treatment for OUD. This aligns with growing interest in various other psychedelic-compounds as potential therapeutics for SUDs and other mental disorders [60, 87, 88].

The long-lasting impact of psychedelics in humans further illustrates the importance of determining the sustained impact of these drugs on VTA function, as even a single dose can create behavioral changes lasting a month or more [89, 90]. The long-term and fast-acting efficacy of these drugs represent a major advantage of psychedelic treatment. Additionally, the expression of the 5-HT2A receptor in the mesolimbic system of human brain [91] supports rodent-based research in the VTA as a primary target for investigating the mechanisms underlying the effects seen in humans. Lastly, a key implication of this research is the potential for co-administering psychedelic microdoses along with addictive drugs such as morphine to prevent the development of dependence from even occurring, rather than solely treating dependence after it has already developed. Our study provides provocative novel cellular evidence to corroborate with human studies, highlighting the need for further examination of psychedelics as treatment agents for substance use disorders such as OUD.

## Acknowledgements

National Institute of Drug Abuse grants R15DA038092 and R15DA049260 from the National Institutes of Health supported this work. The content is solely the responsibility of the authors and does not necessarily represent the official views of the National Institute of Drug Abuse or the National Institutes of Health. This work was also supported by Institutional Mentoring Grants (JE). We thank the NIDA Drug Supply Program for providing the Morphine and LSD used in this study.

## Author contributions

MV and JE designed the experiments. MV acquired and analyzed the electrophysiological data. MV and TR acquired and analyzed behavioral data. TJ, IS and SP acquired and analyzed methylation data. MV, JE and IS wrote the manuscript. All authors contributed to the article and approved the submitted version.

